# Kinetochores promote timely spindle bipolarization to prevent microtubule attachment errors in oocytes

**DOI:** 10.1101/2024.05.20.595072

**Authors:** Shuhei Yoshida, Reiko Nakagawa, Tomoya S. Kitajima

## Abstract

Incorrect kinetochore-microtubule attachment leads to chromosome segregation errors. The risk of incorrect attachments is particularly high in acentrosomal oocytes, where kinetochores are surrounded by randomly oriented microtubules until spindle bipolarization. How the timing of acentrosomal spindle bipolarization is coordinated with kinetochore-microtubule attachment is unknown. Here, we show that in mouse oocytes, kinetochores promote spindle bipolarization when microtubule attachment is unstable in early prometaphase. The kinetochore kinase MPS1 allows this function by regulating multiple pathways, including those mediated by the kinetochore proteins NDC80-NUF2 at their C-terminal domains and the antiparallel microtubule crosslinker PRC1. Inhibition of MPS1 delays spindle bipolarization to late prometaphase, thereby increasing incorrect kinetochore-microtubule attachments. Artificially accelerating spindle bipolarization by overexpressing KIFC1 prevents incorrect kinetochore-microtubule attachment in MPS1-inhibited oocytes. Thus, kinetochores autonomously promote timely acentrosomal spindle bipolarization prior to stabilizing microtubule attachment, thereby reducing the risk of egg aneuploidy.

## Introduction

The spindle is a microtubule-based machine that drives chromosome segregation. In somatic cells, the bipolarity of the spindle is defined by the two centrosomes that act as major microtubule-organizing centers. In contrast, in mammalian oocytes, which do not have canonical centrosomes, microtubules self-assemble a bipolar spindle with no predefined bipolar cue ^1,2^. Spindle assembly starts with microtubule polymerization mainly from cytoplasmic acentriolar microtubule-organizing centers in mouse oocytes ^3^ and from kinetochores in human oocytes ^4,5^, both depending on RanGTP activity elevated around chromosomes ^3,4,6^. The microtubules initially form an apolar ball-shaped spindle, which then transforms into an elongated barrel-shaped spindle through a process called spindle bipolarization. Spindle bipolarization requires the antiparallel microtubule motor KIF11 and is facilitated by numerous microtubule regulators, such as HURP, NuMA, and KIFC1/HSET ^7–10^. Many of these factors are positively regulated by the CDK1 activity ^11–13^, which gradually elevates through prometaphase and metaphase in oocytes ^13,14^. The CDK1 activity also promotes the stability of kinetochore-microtubule attachment ^13^, allowing its gradual increase from late prometaphase to the end of metaphase ^13,15,16^. Thus, the CDK1 activity serves as a master timer that allows the simultaneous progression of spindle bipolarization and kinetochore-microtubule attachment stabilization. However, whether and how these two processes are temporally coordinated remain unknown.

The temporal relationship between spindle bipolarization and kinetochore-microtubule attachment is critical for the fidelity of chromosome segregation. Kinetochores can attach to microtubules prior to spindle bipolarization, but most of such early attempts fail to properly attach the kinetochore pair of the chromosome to the opposite poles of the future bipolar spindle. These early attachments are unstable and undergo error correction after spindle bipolarization, which works efficiently and ensures faithful chromosome segregation in normal somatic cells ^17^. In oocytes, however, early kinetochore-microtubule attachments prior to spindle bipolarization have been implicated as a prevalent contributor to chromosome segregation errors for the following reasons ^1,2,18,19^. First, due to the acentrosomal nature of oocytes, microtubules are randomly oriented around kinetochores prior to spindle bipolarization, increasing the likelihood of erroneous initial attachment ^3,4,15^. Second, error correction of attachments is likely less efficient in oocytes compared to somatic cells due to the predominant regulation of attachment stabilization by CDK1 activity ^13^, which lacks specificity for correct attachments ^16^. Third, the spindle checkpoint, a mechanism that prevents anaphase entry until correct attachments are established, is less stringent in oocytes due to their large cytoplasmic size ^20–22^. Consistent with these notions, in human oocytes, a delay or instability of spindle bipolarization correlates with a subsequent increase in incorrect kinetochore-microtubule attachments ^4^. Accordingly, artificial acceleration of spindle bipolarization may reduce chromosome segregation errors ^10^. However, whether oocytes possess an intrinsic mechanism to prioritize spindle bipolarization before kinetochore-microtubule attachment remains unknown.

Recent reports provide evidence for a functional link between kinetochores and spindle bipolarization in oocytes. In mouse oocytes, NDC80, which forms a heterodimer with NUF2 and serves as a major microtubule anchor for attachment to kinetochores ^23,24^, is essential for spindle bipolarization during meiosis I ^25^. NDC80-NUF2 recruits the antiparallel microtubule crosslinker PRC1 to kinetochores, which promotes KIF11-mediated spindle bipolarization ^25^. In human oocytes, kinetochore localization of PRC1 is not detected ^25^, but instead oocyte-specific microtubule organizing centers localize to kinetochores ^5^. These observations are consistent with the idea that kinetochores provide a common platform for promoting acentrosomal spindle assembly via divergent molecular mechanisms among mammalian species. Identification of upstream regulators of kinetochores may contribute to the molecular understanding of a conserved regulatory principle of acentrosomal spindle assembly in mammals.

In this study, we find that in mouse oocytes, MPS1, a kinase active at unattached kinetochores ^26,27^, promotes timely spindle bipolarization. MPS1 exerts this function via NDC80-NUF2 at their C-terminal domains and PRC1. Inhibition of MPS1 delays spindle bipolarization, which causes chromosome misalignment with incorrect kinetochore-microtubule attachments. We propose that kinetochores act as a platform to timely initiate spindle bipolarization prior to stabilizing microtubule attachment, thereby reducing the risk of chromosome segregation errors in acentrosomal oocytes.

## Results

### MPS1 activity is required for acentrosomal spindle bipolarization in the absence of stable kinetochore-microtubule attachment

In mouse oocytes, kinetochore NDC80-NUF2 plays a dual role in establishing microtubule attachment and in promoting acentrosomal spindle bipolarization ^25^. NDC80-NUF2 can promote acentrosomal spindle bipolarization without stable kinetochore-microtubule attachment ^25,28^. One of the candidate factors that regulate kinetochore-based spindle bipolarization is MPS1, a kinase active at kinetochores lacking stable microtubule attachment ^26,27^. To test spindle bipolarization independent of stable kinetochore-microtubule attachment, we replaced endogenous NDC80 with NDC80-9D, a phospho-mimetic mutant form deficient in stabilizing kinetochore-microtubule attachment ^23,24^, by deleting the floxed *Ndc80* gene with the oocyte-specific *Zp3*-Cre recombinase and exogenously expressing NDC80-9D through microinjection into oocytes ^25,28^. Oocytes were monitored for spindle formation with the microtubule marker EGFP-MAP4 and the chromosome marker H2B-mCherry by live confocal microscopy ^3^. We analyzed the dynamics of spindle morphology by measuring the sphericity and aspect ratio of an ellipsoid fitted to the 3D mass of microtubule signals (Fig. 1A). This analysis showed that NDC80-9D-expressing oocytes initiated spindle bipolarization with a kinetics similar to wild-type NDC80 (NDC80-WT)-expressing oocytes, although it resulted in an excessively elongated bipolar spindle (Fig. 1B), confirming our previous report ^28^. We then treated NDC80-9D-expressing oocytes with reversine, a specific inhibitor of MPS1 ^29^. Interestingly, we found that MPS1-inhibited, NDC80-9D-expressing oocytes exhibited an irregularly shaped spindle that did not fit well with an ellipsoid, as indicated by its low sphericity, throughout meiosis I (Fig. 1C). These observations suggest that MPS1 inhibition severely perturbs spindle bipolarization in the absence of stable kinetochore-microtubule attachment. In contrast, the effect of MPS1 inhibition was much less pronounced in NDC80-WT-expressing oocytes (Fig. 1C). MPS1-inhibited NDC80-9D-expressing oocytes, but not NDC80-9A-expressing oocytes, failed to form a bipolar spindle even when arrested for a prolonged period at metaphase I with proTAME, an inhibitor of the anaphase promoting complex ^30^ (Fig. S1A, B). In contrast, during meiosis II, MPS1 inhibition only modestly prevented spindle bipolarization in NDC80-9D-expressing oocytes (Fig. 1C), consistent with previous reports that kinetochore-independent pathways support spindle bipolarization during meiosis II ^25,31^. We noticed that MPS1 inhibition reduced NDC80-9D levels at kinetochores to ∼73% at early prometaphase (Fig. S1C), suggesting that MPS1 promotes kinetochore localization of phosphorylated NDC80. Consistently, kinetochore NDC80 levels just after M-phase entry (1 hour after nuclear envelope breakdown, NEBD) were significantly reduced by MPS1 inhibition, especially in oocytes treated with nocodazole, a microtubule depolymerizing drug (Fig. S1D). However, the reduced localization of NDC80-9D in MPS1-inhibited oocytes was unlikely to sufficiently explain their severe spindle defects, because lower expression of NDC80-9D, which resulted in its kinetochore levels comparable to those of MPS1-inhibited NDC80-9D (∼56%, Fig. S1E), did not recapitulate the severe spindle defects (Figure S1F). These results suggest that MPS1 kinase activity is essential for acentrosomal spindle bipolarization in the absence of stable kinetochore-microtubule attachment by mechanisms in addition to facilitating NDC80 kinetochore localization.

**Figure 1.**
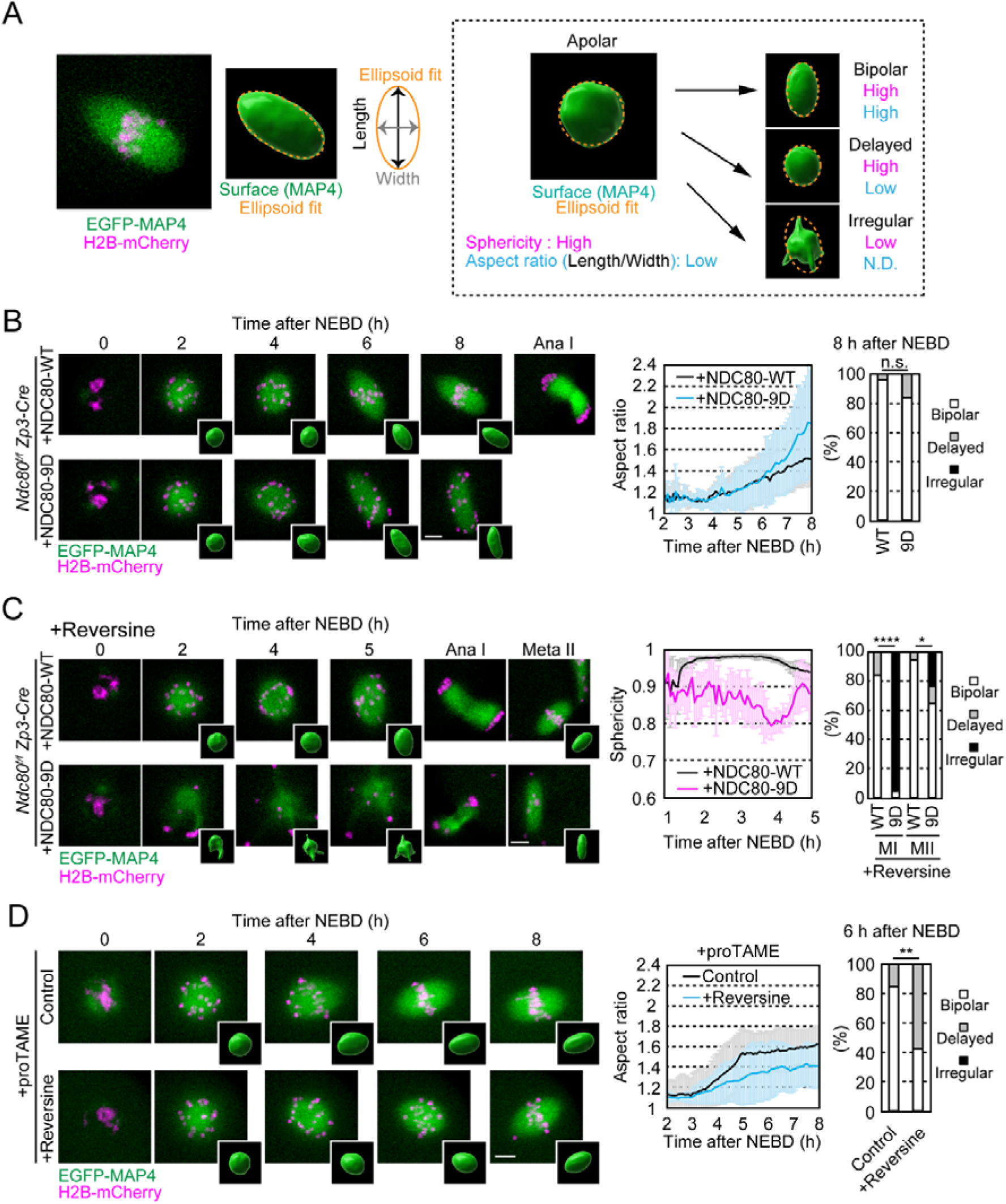
MPS1 activity is required for acentrosomal spindle bipolarization in the absence of stable kinetochore-microtubule attachment. **A.** Spindle morphology quantification. Representative z-projection image of EGFP-MAP4 (spindle, green) and H2B-mCherry (chromosome, magenta) at metaphase I in the mouse oocyte. The surface of 3D reconstructed EGFP-MAP4 is fitted to an ellipsoid. Examples shown are identical to those in B and C. Based on the aspect ratio of the ellipsoid, spindles are classified as “bipolar” or “delayed”. Spindles that did not fit well with an ellipsoid are classified as “irregular” (see Methods). **B.** Spindle bipolarization in NDC80-9D oocytes. Live imaging of *Ndc80^f/f^ Zp3-Cre* oocytes expressing NDC80-WT/-9D. Insets show 3D reconstructed spindles. Temporal changes in the aspect ratio of the spindle (mean ± SD, n=6, 6 oocytes) and morphology classification at 8 hours after nuclear envelope breakdown (NEBD) are shown (n=18, 17 oocytes from 3 independent experiments). n.s., not significant by Fisher’s exact test for “bipolar” groups. **C.** MPS1 inhibition impairs spindle bipolarization in NDC80-9D oocytes. Live imaging of *Ndc80^f/f^ Zp3-Cre* oocytes expressing NDC80-WT/-9D with the MPS1 inhibitor reversine. Temporal changes in the sphericity of the spindle (mean ± SD, n=6, 6 oocytes) and morphology classification at 5 hours after NEBD (meiosis I, MI) and MII (meiosis II) (n=25, 23, 18, 17 oocytes from 3 independent experiments) are shown. *p=0.0408, ***p=0.0001, and ****p<0.0001 by Fisher’s exact test for “bipolar” groups. **D.** MPS1 inhibition delays spindle bipolarization. Live imaging of oocytes with proTAME, an inhibitor of the anaphase promoting complex. Temporal changes in the aspect ratio of the spindle (mean ± SD, n=7, 7 oocytes) and morphology classification at 6 hours after NEBD (n=26, 26 from 4 independent experiments) are shown. **p=0.0034 by Fisher’s exact test for “bipolar” groups. Scale bars, 10 μm.

### MPS1 is required for initiating spindle bipolarization during early prometaphase

In oocytes, stable kinetochore-microtubule attachment is rarely observed during prometaphase and gradually increases during metaphase ^13,15,16^. Therefore, if MPS1 promotes spindle bipolarization in the absence of stable kinetochore-microtubule attachment, inhibition of MPS1 should lead to a delay in spindle bipolarization during prometaphase. However, such a delay has not been reported in previous studies ^32,33^ or noticeable in our data (Fig. 1C). We speculated that because MPS1 inhibition compromises the spindle checkpoint ^26,32^ and thereby accelerates the onset of anaphase spindle elongation, these effects may have masked a spindle bipolarization phenotype during prometaphase and metaphase. Therefore, we tested MPS1 inhibition in oocytes treated with proTAME, which blocks anaphase entry ^30^. Under the proTAME-treated condition, we found that MPS1 inhibition significantly delayed spindle bipolarization (Fig. 1D). Thus, MPS1 is required for initiating spindle bipolarization during early prometaphase, when kinetochore-microtubule attachment is unstable.

### MPS1 promotes spindle bipolarization via the C-terminal domains of NDC80-NUF2

We investigated how MPS1 activity promotes spindle bipolarization. The C-terminal domains of NDC80-NUF2 are not directly involved in microtubule attachment but promote spindle bipolarization ^25^. To specifically test whether the function of the C-terminal domains of NDC80-NUF2 in spindle bipolarization depends on MPS1 activity, we used the C-terminal fragments of NDC80 (a.a. 461–642, termed NDC80ΔN) and NUF2 (a.a. 276–463, termed NUF2ΔN) (Fig. 2A), which localize to kinetochores and partially rescue spindle bipolarization when co-expressed in *Ndc80*-deleted oocytes ^25^. Notably, MPS1 inhibition largely diminished the kinetochore localization of NDC80ΔN (Fig. S2A), consistent with the idea that MPS1 contributes to NDC80 localization at kinetochores, and severely perturbed spindle bipolarization (Fig. 2B).

**Figure 2.**
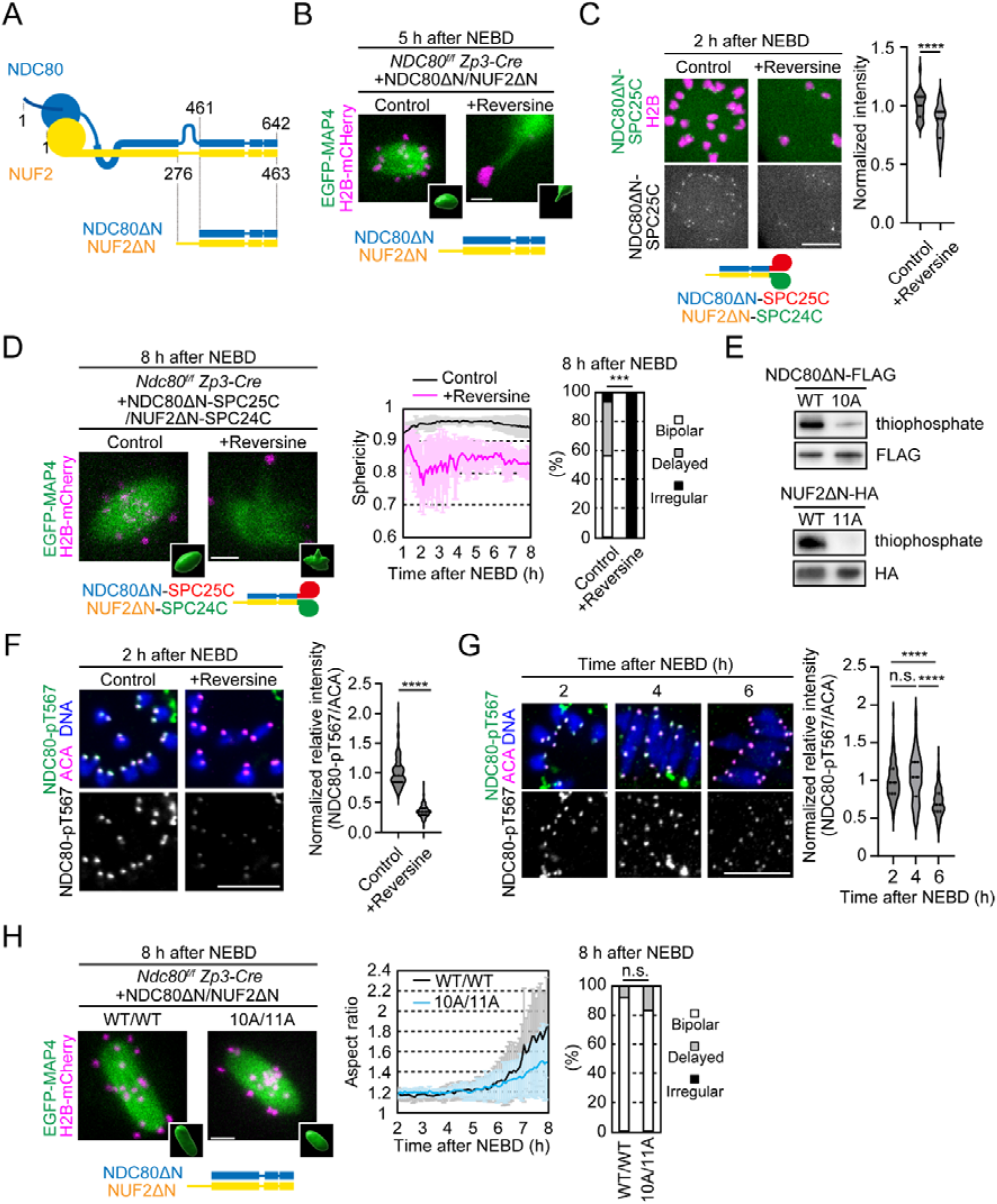
MPS1 promotes spindle bipolarization via the C-terminal domains of NDC80-NUF2 during prometaphase. **A.** Diagram of NDC80 and NUF2. **B.** The C-terminal domains of NDC80 and NUF2 require MPS1 for spindle bipolarization. Live imaging of *Ndc80^f/f^ Zp3-Cre* oocytes expressing EGFP-MAP4 (spindle, green), H2B-mCherry (chromosome, magenta), NDC80ΔN, and NUF2ΔN, treated with reversine. Z-projection and 3D-reconstruction images are shown. **C.** Tethering the C-terminal domains of NDC80 and NUF2 at kinetochores. Live imaging of *Ndc80^f/f^ Zp3-Cre* oocytes expressing NDC80ΔN-SPC25C-mNeonGreen, NUF2ΔN-SPC24C, and H2B-mCherry, treated with reversine. SPC25C (a.a. 120–226) and SPC24C (a.a. 122–201) are kinetochore-targeting domains. Normalized intensities of NDC80ΔN-SPC25C-mNeonGreen are shown (median and quartiles, n=30, 30 kinetochores of 3 oocytes from 1 experiment). ****p<0.0001 by two-tailed unpaired Student’s t-test. **D.** The C-terminal domains of NDC80 and NUF2 tethered at kinetochores require MPS1 for spindle bipolarization. Live imaging of *Ndc80^f/f^ Zp3-Cre* oocytes expressing EGFP-MAP4 (spindle, green), H2B-mCherry (chromosome, magenta), NDC80ΔN-SPC25C and NUF2ΔN-SPC24C, treated with reversine, in the presence of proTAME. Temporal changes in the sphericity of the spindle (mean ± SD, n=16, 16 oocytes) and morphology classification at 8 hours after NEBD (n=16, 16 oocytes) are shown. ***p=0.0008 by Fisher’s exact test for “bipolar” groups. **E.** Phosphorylation of NDC80ΔN and NUF2ΔN by MPS1 *in vitro*. Western blotting of NDC80-WT/-10A-FLAG and NUF2-WT/-11A-HA. *In vitro* phosphorylation was performed with ATPγS, which was detected by Western blotting against thiophosphate. **F.** MPS1-dependent NDC80 phosphorylation. Immunostaining with anti-phosphorylated NDC80 at T567 (NDC80-pT567), ACA (kinetochores), and Hoechst33342 (DNA). Normalized relative intensities of NDC80-pT567 are shown (median and quartiles, n=200, 200 kinetochores from 5, 5 oocytes. Three independent experiments were performed). ****p<0.0001 by two-tailed unpaired Student’s t-test. **G.** NDC80-T567 phosphorylation during meiosis I. Immunostaining or oocytes with anti-NDC80-pT567, ACA (kinetochores), and Hoechst33342 (DNA). Normalized relative intensities of NDC80-pT567 are shown (median and quartiles, n=200, 199, 200 kinetochores from 5, 5, 5 oocytes. Three independent experiments were performed). n.s., not significant, ****p<0.0001 by Tukey’s multiple comparison test. **H.** Phospho-mutants of NDC80ΔN and NUF2ΔN do not recapitulate MPS1 inhibition. Live imaging of *Ndc80^f/f^ Zp3-Cre* oocytes expressing EGFP-MAP4 (spindle, green), H2B-mCherry (chromosome, magenta), NDC80ΔN-WT/-10A and NUF2ΔN-WT/-11A. Temporal changes in the aspect ratio of the 3D reconstructed spindle (mean ± SD, n=10, 12 oocytes) and morphology classification at 8 hours after NEBD are shown (n=28, 26 oocytes from 3 independent experiments). n.s., not significant by Fisher’s exact test for “bipolar” groups. Scale bars, 10 μm.

To test additional contributions of MPS1 to spindle bipolarization via the C-terminal domains of NDC80-NUF2, we tethered NDC80ΔN and NUF2ΔN to kinetochores by fusing them with the kinetochore-targeting domains of SPC25 and SPC24 (NDC80ΔN-SPC25C and NUF2ΔN-SPC24C), respectively ^25^. As expected, a substantial amount of NDC80ΔN-SPC25C (∼84%) was retained at kinetochores after MPS1 inhibition (Fig. 2C). Nevertheless, MPS1 inhibition severely impaired the ability of NDC80ΔN-SPC25C and NUF2ΔN-SPC24C to rescue spindle bipolarization, resulting in the formation of an irregularly shaped spindle that did not fit well with an ellipsoid throughout meiosis I (Fig. 2D). These results suggest that MPS1 promotes spindle bipolarization via the C-terminal domains of NDC80-NUF2 at kinetochores, in addition to ensuring NDC80-NUF2 localization.

### MPS1 directly phosphorylates the C-terminal domains of NDC80-NUF2 during prometaphase

We searched for MPS1-mediated phosphorylation sites on the C-terminal domains of NDC80 and NUF2. *In vitro* kinase assay using recombinant NDC80ΔN, NUF2ΔN and MPS1 followed by mass spectrometry analysis identified 21 candidate phosphorylation sites on the C-terminal domains of NDC80 and NUF2 (NDC80-T485, T491, T494, S496, T499, S544, T567, T572, S595, and S616; and NUF2-S311, S312, S329, T336, S340, T354, T392, S403, S419, S445, and T452) (Fig. 2E). We produced phospho-specific antibodies against phosphorylated NDC80-T567, which detected kinetochores in oocytes in a manner dependent on MPS1 activity and NDC80 (Fig. 2F and S2B). Substitution of NDC80-T567 with alanine substantially reduced the phospho-antibody signals at kinetochores in oocytes (Fig. S2C), demonstrating the specificity of the antibody. Levels of phosphorylated NDC80-T567 at kinetochores were high during prometaphase and decreased during metaphase (Fig. 2G), consistent with the idea that MPS1 is more active at kinetochores with less stable attachments. These results suggest that MPS1 phosphorylates the C-terminal domains of NDC80 and NUF2 during prometaphase, when spindle bipolarization initiates. However, substitution of all 21 candidate phosphorylation sites on the C-terminal domains of NDC80 and NUF2 (NDC80ΔN-10A and NUF2ΔN-11A) did not significantly reduce their ability to bipolarize the spindle in *Ndc80*-deleted oocytes (Fig. 2H). Thus, additional MPS1 target sites on NDC80 and NUF2, or on other proteins, likely facilitate spindle bipolarization through the C-terminal domains of NDC80-NUF2.

### MPS1 activity promotes kinetochore localization and spindle bipolarization activity of PRC1

One of the pathways downstream of the C-terminal domains of NDC80-NUF2 is the antiparallel microtubule crosslinker PRC1, which is recruited to kinetochores and promotes spindle bipolarization ^25^. Interestingly, we found that MPS1 inhibition greatly reduced the kinetochore levels of PRC1 at early prometaphase (2 hours after NEBD) (Fig. 3A). This reduction was not attributable to reduced NDC80 by MPS1 inhibition, because kinetochore NDC80 levels were not significantly decreased by MPS1 inhibition at this stage (Fig. S3A). These results suggest that MPS1 promotes the recruitment of PRC1 to kinetochores.

**Figure 3.**
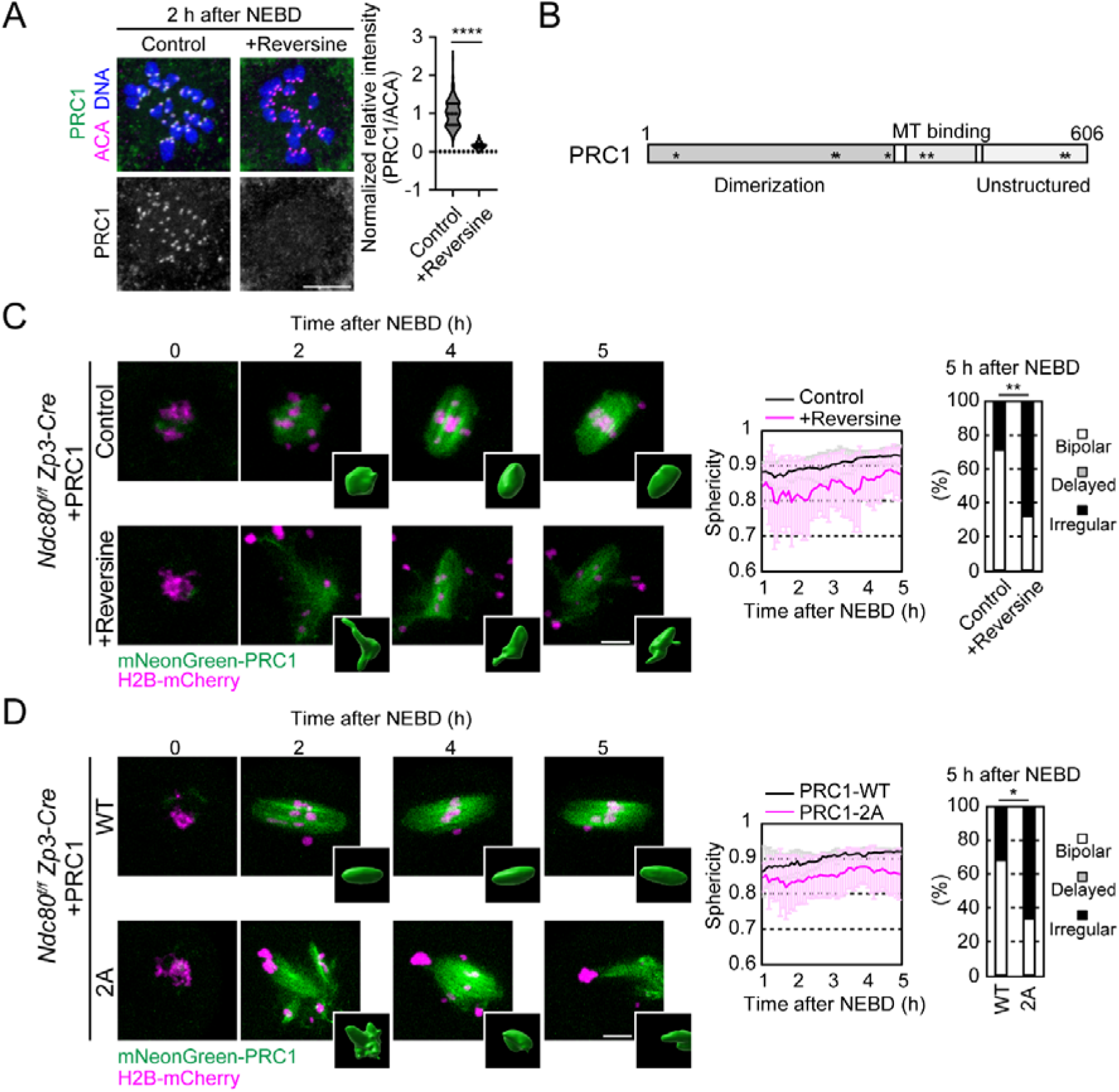
MPS1 activity promotes kinetochore localization and spindle bipolarization activity of PRC1. **A.** MPS1 promotes kinetochore localization of PRC1. Immunostaining with anti-PRC1, ACA (kinetochores), and Hoechst33342 (DNA). Normalized relative intensities are shown (median and quartiles, n=240, 240 kinetochores from 6, 6 oocytes. Three independent experiments were performed). ****p<0.0001 by two-tailed unpaired Student’s t-test. **B.** Phosphorylated amino acids of PRC1 by MPS1 kinase *in vitro* are shown. **C.** PRC1 activity for spindle bipolarization depends on MPS1. Live imaging of *Ndc80^f/f^ Zp3-Cre* oocytes expressing mNeonGreen-PRC1 (green) and H2B-mCherry (chromosome, magenta), treated with reversine. Insets show 3D reconstructed images. Temporal changes in the sphericity of the spindle (mean ± SD, n=10, 10 oocytes) and morphology classification at 5 hours after NEBD are shown (n=28, 28 oocytes from 3 independent experiments). **p=0.0069 by Fisher’s exact test for “bipolar” groups. **D.** Two potential phosphorylation sites on PRC1 are critical for its spindle bipolarization activity. As in C, mNeonGreen-PRC1-WT/-2A (T578 and S583 substituted to alanine)-expressing oocytes were tested (sphericity, mean ± SD, n=6, 6 oocytes; morphology classification, n=22, 24 oocytes from 3 independent experiments, *p=0.0377 by Fisher’s exact test for “bipolar” groups). Scale bars, 10 μm.

Consistent with the retained ability of NDC80ΔN-10A and NUF2ΔN-11A to promote spindle bipolarization, they recruited PRC1 to kinetochores in *Ndc80*-deleted oocytes, similarly to NDC80ΔN and NUF2ΔN (Fig. S3B). We therefore speculated that PRC1 is also a direct target of MPS1, in addition to NDC80-NUF2. Mass spectrometry analysis of recombinant PRC1 phosphorylated by MPS1 *in vitro* detected T40, T265, S267, T327, T379, T398, T578, and S583 as candidate phosphorylation sites (Fig. 3B). Substitution of all 8 candidate phosphorylation sites (PRC1-8A) did not largely affect the ability of PRC1 to localize to kinetochores (Fig. S3C), indicating that these phosphorylation sites are not essential for kinetochore localization of PRC1. We then explored the possibility that MPS1-mediated phosphorylation on these sites regulates PRC1 activity for spindle bipolarization independently of regulating its kinetochore localization. To evaluate the ability of PRC1 to promote spindle bipolarization independently of its kinetochore localization, we used *Ndc80*-deleted oocytes, where overexpression of PRC1 rescues spindle bipolarization defects without its kinetochore localization ^25^. We found that MPS1 inhibition significantly prevented overexpressed PRC1 from rescuing spindle bipolarization defects in *Ndc80*-deleted oocytes (Fig. 3C). Notably, we found that PRC1-2A, which carries alanine substitutions at two of the candidate phosphorylation sites (T578 and S583) in the C-terminal unstructured domain, largely recapitulated the failure of rescue when expressed in *Ndc80*-deleted oocytes, while retaining its ability to localize to spindle microtubules (Fig. 3D). These results suggest that MPS1 directly regulates PRC1 activity to promote spindle bipolarization.

### MPS1-mediated timely spindle bipolarization prevents kinetochore-microtubule attachment errors

Our results demonstrate that MPS1 promotes spindle bipolarization via multiple pathways independent of kinetochore-microtubule attachment during early prometaphase. However, MPS1 is not essential for spindle bipolarization because attachment-dependent pathways can support spindle bipolarization with a delay to late prometaphase in MPS1-inhibited oocytes (Fig. 1C). These findings led us to ask the significance of the initiation of spindle bipolarization during early prometaphase. In MPS1-inhibited oocytes, the delay in spindle bipolarization was accompanied by a significant increase in misaligned chromosomes (Fig. 4A). The chromosomes showed a significant increase in incorrect attachment of kinetochores with cold-stable microtubules at early metaphase (Fig. 4B). Thus, MPS1 promotes spindle bipolarization and prevents kinetochores from microtubule attachment errors.

**Figure 4.**
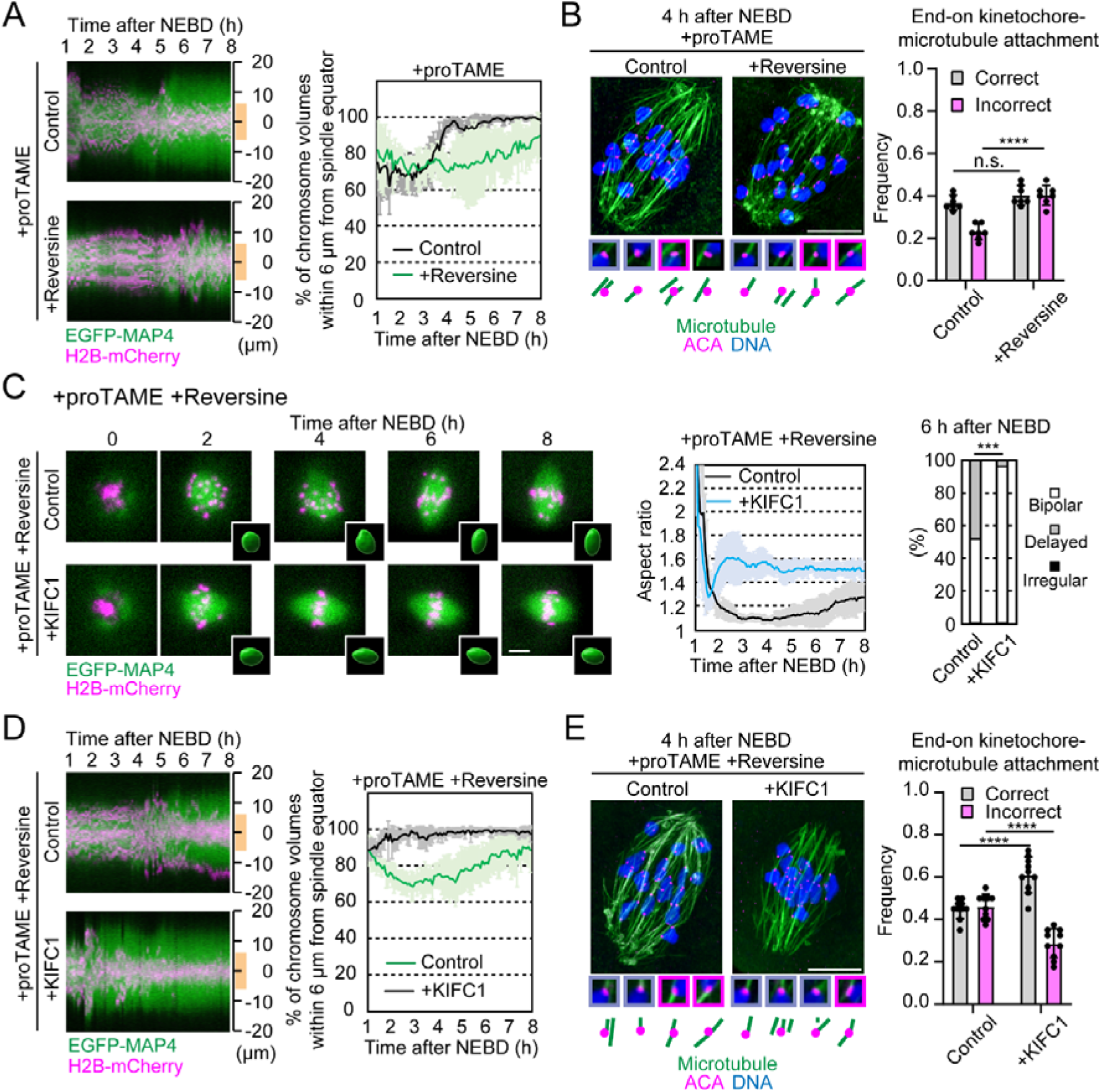
MPS1-mediated timely spindle bipolarization prevents kinetochore-microtubule attachment errors. **A.** MPS1 inhibition causes chromosome misalignment. Images of EGFP-MAP4 (spindle, green) and H2B-mCherry (chromosome, magenta) acquired in the experiments in Fig. 1D were used to generate kymographs along the spindle axis in 3D. Distance from the spindle equator is shown on the right. To analyze chromosome alignment, the percentage of chromosome volume within 6 μm from the spindle equator (orange line) was calculated (mean ± SD, n=7, 7 oocytes). **B.** MPS1 inhibition causes kinetochore-microtubule attachment errors. After brief cold treatment, oocytes were fixed and immunostained for stable microtubules (green), kinetochores (magenta), and DNA (blue). Magnified images of end-on monopolar (correct, gray frame) and merotelic (incorrect, magenta frame) attachments are shown (mean ± SD, n=7, 7 oocytes from 3 independent experiments). n.s., not significant, ****p<0.0001 by two-tailed unpaired Student’s t-test. **C.** KIFC1 overexpression accelerates spindle bipolarization in MPS1-inhibited oocytes. Live imaging of oocytes expressing EGFP-MAP4 (spindle, green), H2B-mCherry (chromosome, magenta), and KIFC1, treated with reversine and proTAME. Temporal changes in the aspect ratio of the spindle (mean ± SD, n=9, 9 oocytes) and morphology classification at 6 hours after NEBD are shown (n=29, 27 oocytes from 3 independent experiments). ***p=0.0002 by Fisher’s exact test for “bipolar” groups. **D.** KIFC1 overexpression rescues chromosome misalignment in MPS1-inhibited oocytes. Oocyte images acquired in the experiment shown in C were analyzed for chromosome alignment as in A (mean ± SD, n=9, 9 oocytes). **E.** KIFC1 overexpression prevents kinetochore-microtubule attachment errors in MPS1-inhibited oocytes. Kinetochore-microtubule attachments in KIFC1-expressing oocytes treated with reversine and proTAME were analyzed as in B (mean ± SD, n=10, 10, 10, 10 oocytes from 4 independent experiments). ****p<0.0001 by two-tailed unpaired Student’s t-test. Scale bars, 10 μm.

We hypothesized that in MPS1-inhibited oocytes, the delay in spindle bipolarization caused chromosome misalignment with incorrect kinetochore-microtubule attachment. If this holds true, artificial acceleration of spindle bipolarization should facilitate chromosome alignment with correct kinetochore-microtubule attachment in MPS1-inhibited oocytes. To test this idea, we used overexpression of KIFC1/HSET, a microtubule motor that accelerates spindle bipolarization ^9,10^. As expected, KIFC1 overexpression significantly accelerated spindle bipolarization in MPS1-inhibited oocytes (Fig. 4C). Remarkably, these oocytes exhibited significantly rescued chromosome alignment (Fig. 4D) with a significant decrease in incorrect attachment and increase in correct attachment of kinetochores with cold-stable microtubules (Fig. 4E). These results suggest that MPS1 activity prevents incorrect kinetochore-microtubule attachment by timely initiating spindle bipolarization during prometaphase.

## Discussion

In oocytes, due to the absence of centrosomes, kinetochores are surrounded by randomly oriented microtubules during early prometaphase prior to spindle bipolarization. This situation favors kinetochores to form improper microtubule attachments, which can lead to chromosome segregation errors ^1,2,18,19^. In this study, we identified the MPS1 kinase as a key player that allows kinetochores to actively promote spindle bipolarization before stabilizing their microtubule attachments. MPS1 kinase activity is essential for spindle bipolarization when kinetochore-microtubule attachment is unstable, by regulating multiple downstream factors, including the C-terminal regions of NDC80 and NUF2, and PRC1. Oocytes lacking MPS1 kinase activity exhibit delayed spindle bipolarization, which causes increased incorrect kinetochore-microtubule attachment. Thus, kinetochores are not only passively driven by the spindle, but rather actively promote spindle bipolarization before stabilizing their own microtubule attachment, thereby facilitating correct chromosome segregation.

Based on these findings, we propose that acentrosomal spindle bipolarization is driven by two distinct modes that act sequentially on kinetochores during meiosis I (Fig. 5). The first mode is mediated by kinetochores with unstable microtubule attachment. In this mode, kinetochores employ MPS1 activity to create a microenvironment that concentrates microtubule regulators, such as the antiparallel microtubule crosslinker PRC1, via NDC80-NUF2 ^25^, which facilitates KIF11-mediated bipolar microtubule sorting and thereby initiates spindle bipolarization during early prometaphase. The first mode leads to the acquisition of spindle bipolarity, which increases the likelihood of subsequent microtubule attachments of the kinetochore pair of chromosomes to both spindle poles. Kinetochores drive the first mode until the gradual increase in CDK1 activity stabilizes microtubule attachment ^13,16^, which reduces MPS1 activity at kinetochores ^34,35^, allowing kinetochores to switch to the second mode. In the second mode, kinetochores employ stably attached microtubules, which maintains the integrity of the bipolar spindle by restricting microtubule organizing centers at the poles and preventing excessive spindle elongation during late prometaphase and metaphase ^28^. In oocytes lacking MPS1 activity, the first mode is absent and thus spindle bipolarization is delayed, while increased CDK1 activity stabilizes microtubules attached to kinetochores from random directions. Subsequently, the second mode initiates with kinetochores stably attached by microtubules, which mediate spindle bipolarization and thereby remain as incorrect, merotelic attachments in the resulting bipolar spindle. According to this model, in normal oocytes, kinetochores autonomously coordinate the temporally sequential action of the two distinct modes for bipolar spindle assembly to prevent the formation of incorrect kinetochore-microtubule attachments.

**Figure 5.**
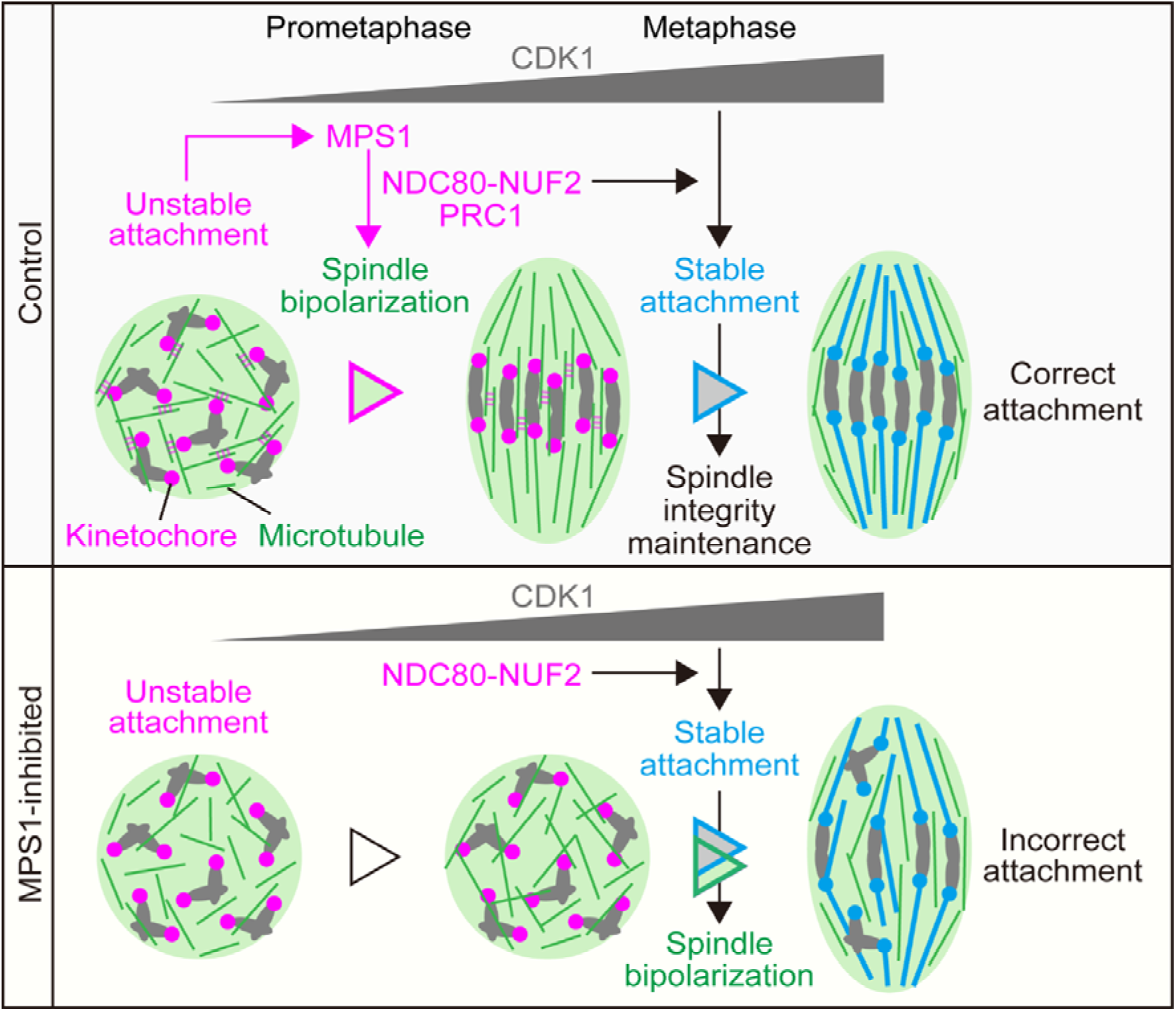
Kinetochores promote timely spindle bipolarization to prevent microtubule attachment errors in oocytes. During prometaphase, kinetochores with unstable microtubule attachment employ MPS1 to promote spindle bipolarization via the C-terminal regions of NDC80-NUF2, and PRC1. Spindle bipolarity facilitates bipolar microtubule attachment to the kinetochore pair on chromosomes. In parallel, the gradual increase in CDK1 activity stabilizes kinetochore-microtubule attachment. The stably attached microtubules contribute to the maintenance of bipolar spindle integrity during metaphase. However, in oocytes lacking MPS1 activity, spindle bipolarization is delayed, while CDK1 activity gradually increases, which stabilizes kinetochore-microtubule attachment. The microtubules stably attached to kinetochores participate in spindle bipolarization and thus remain as merotelic microtubules in the bipolar spindle.

Our results show that one of the critical roles of MPS1 kinase in oocytes is the timely initiation of spindle bipolarization, in addition to spindle checkpoint activation and centromeric cohesion protection ^32,33^. In somatic cells, chromosome misalignment induced by MPS1 inhibition is attributed to loss of MPS1-mediated direct regulation of kinetochore-microtubule attachment ^36–38^. In contrast, in oocytes, a substantial fraction of misaligned chromosomes induced by MPS1 inhibition are attributed to delayed spindle bipolarization, as chromosome misalignment and incorrect kinetochore-microtubule attachment in MPS1-inhibited oocytes are largely suppressed by artificially accelerating spindle bipolarization. We suggest that the MPS1-mediated timely initiation of spindle bipolarization is particularly critical for acentrosomal oocytes.

MPS1 kinase activity promotes spindle bipolarization likely through multiple pathways in oocytes. First, MPS1 ensures proper localization of NDC80 to kinetochores. Second, MPS1 regulates the C-terminal regions of NDC80 and NUF2. Our data show that MPS1 directly phosphorylates these regions in oocytes, although it is unclear whether direct phosphorylation contributes to timely spindle bipolarization. Third, MPS1 regulates PRC1, an antiparallel microtubule crosslinker that promotes spindle bipolarization ^25,39,40^. MPS1 promotes kinetochore localization and spindle bipolarizing activity of PRC1. Our data do not exclude the possibility that MPS1 has additional downstream pathways to promote spindle bipolarization. Although future studies are needed to fully understand the downstream molecular pathways of MPS1, our current results indicate that MPS1 is a master kinase that controls multiple pathways to promote timely spindle bipolarization at kinetochores prior to stabilization of microtubule attachment.

Kinetochores contribute to bipolar spindle assembly through distinct mechanisms in mouse and human oocytes. In mouse oocytes, kinetochores with unstable microtubule attachment employ MPS1 to promote spindle bipolarization, whereas in human oocytes, kinetochores recruit microtubule organizing centers to promote microtubule polymerization ^5^. Whether human oocytes require MPS1 kinase activity for MTOC recruitment, microtubule polymerization, or spindle bipolarization is an intriguing question for future studies.

In summary, this study found a kinetochore-based strategy principle in acentrosomal oocytes that prevents the risk of chromosome segregation errors by actively promoting spindle bipolarization before stabilizing microtubule attachment. This finding provides a basis for future research to develop approaches to prevent egg aneuploidy, a major cause of pregnancy loss and congenital disease.

## Materials and methods

### Mice

All animal experiments were approved by the Institutional Animal Care and Use Committee of RIKEN Kobe Branch (IACUC). B6D2F1 (C57BL/6 x DBA/2), *Ndc80^flox/flox^ Zp3-Cre* female mice ^25^, 8–16 weeks old, were used to obtain oocytes.

### Mouse oocyte culture

Mice were injected with 0.1 ml of CARD HyperOva (KYUDO). Fully grown oocytes were collected 48 hours after injection and placed in M2 medium containing 200 μM 3-isobutyl-1-methyl-xanthine (IBMX, Sigma) at 37°C. Meiotic resumption was induced by washing to remove IBMX. When indicated, 1 μM reversine (Cayman), 5 μM proTAME (RD systems), 66 μM nocodazole (Sigma) were used. DMSO was used as a control.

### mRNA synthesis and injection

mRNAs were transcribed *in vitro* by using the mMESSAGE mMACHINE T7 kit (Invitrogen). The mRNAs were introduced into fully grown oocytes by microinjection. The microinjected oocytes were cultured at 37°C for 3–4 hours before IBMX washing. Microinjections were performed with mRNA of EGFP-MAP4 (3 pg); H2B-mCherry (0.15 pg); NDC80-WT, −9A, −9D, and T567A (1 pg); NDC80ΔN-WT and −10A (1 pg); NDC80ΔN-mNeonGreen (1 pg); NUF2ΔN-WT and −11A (1 pg); NDC80ΔN-SPC25C (1 pg); NDC80ΔN-SPC25C-mNeonGreen (1 pg); NUF2ΔN-SPC24C (1 pg); mNeonGreen-PRC1-WT and −2A (2 pg); NDC80-9D-GFP (1 pg, 0.1 pg); KIFC1 (0.12 pg); 24xGCN-PRC1-WT, −8A (0.1 pg); and scFv-sfGFP (0.25 pg).

### Live imaging

A customized Zeiss LSM710 or LSM880 confocal microscope equipped with a 40x C-Apochromat 1.2 NA water immersion objective lens (Carl Zeiss) was controlled by Zen software using the multi-position autofocus macros AutofocusScreen ^41^ and MyPic ^42^. For spindle and chromosome imaging, we recorded 11 z-confocal sections (every 4 μm) of 512 x 512 pixel xy images at 5–6 minutes time intervals for at least 12 hours after the induction of meiotic resumption.

### 4D spindle analysis

To analyze spindle morphology, we performed 3D surface rendering of EGFP-MAP4 or mNeonGreen-PRC1 signals using Imaris software (Oxford Instruments). The generated 3D surface was fitted to an ellipsoid, which was used to categorize spindle morphology. If the volume of the fitted ellipsoid was >1.05-fold greater than that of the 3D surface of the spindle, indicating that the spindle did not fit well to an ellipsoid, the spindle was classified as “irregular”. Otherwise, the aspect ratio of the length (longest axis length) to the width (average of two shorter axis lengths) of the fitted ellipsoid and its sphericity were calculated. Spindles with an aspect ratio greater than 1.2 were classified as “bipolar”, and others as “delayed”.

### Immunostaining of oocytes

Oocytes were fixed with 1.6% formaldehyde (methanol-free) in 10 mM PIPES (pH 7.0), 1 mM MgCl_2_, and 0.1% Triton X-100 for 30 minutes at room temperature. To detect cold-stable microtubules, oocytes were incubated in ice-cold M2 medium for 10 min before fixation (cold treatment). After fixation, oocytes were washed and permeabilized with PBT (PBS with 0.1% Triton X-100) at 4°C overnight. After blocking with 3% bovine serum albumin (BSA)-PBT for 1 hour, oocytes were incubated with primary antibodies at 4°C overnight. Oocytes were washed with 3% BSA-PBT and then incubated with secondary antibodies and 20 μg/ml Hoechst33342 for 2 hours for kinetochore imaging or overnight for cold-stable microtubule imaging. Oocytes were washed again and stored in 0.01% BSA-PBS. Oocytes were imaged using a Zeiss LSM780, LSM980 confocal microscope for kinetochore imaging, or a Zeiss LSM880 confocal microscope with AiryScan for cold-stable microtubule imaging.

The following primary antibodies were used: a rabbit anti-NDC80 antiserum (1:2000, a gift from Dr. Robert Benezra), a human anti-centromere protein antibody (ACA, 1:500, 15-234, Antibodies Incorporated), a rat monoclonal anti-alpha tubulin (1:2000, MCA77G, Bio-Rad), a rat anti-GFP antibody (1:500, GF090R 04404-84, Nacalai), and a rabbit anti-PRC1 (1:100, H-70 sc-8356, Santa Cruz). The following secondary antibodies were used: Alexa Fluor 488 goat anti-rabbit IgG (H+L) (A11034), goat anti-rat IgG (H+L) (A11006), and Alexa Fluor 555 goat anti-human IgG (H+L) (A21433) (1:500, Thermo Fisher).

### Phospho-specific antibodies against phosphorylated NDC80-T567

To produce a rabbit polyclonal antibody against NDC80-pT567, the phospho-peptide Cys+QREYQL(pT)VKTTT was synthesized and used for the immunization of a rabbit. The anti-NDC80-pT567 antibody was affinity purified from the immunized serum using the same phospho-peptide. The fraction that binds to the non-phosphorylated peptide Cys+QREYQLTVKTTT was absorbed.

### Quantification of signal intensity

Fiji (https://fiji.sc/) and Imaris software (Oxford Instruments) were used to quantify fluorescence signals. We measured fluorescence intensity for NDC80, PRC1, or GFP at kinetochores and subtracted the signal intensity at a cytoplasmic region near the kinetochore. Similarly, we measured the fluorescence intensity for ACA at the same kinetochores. We calculated the ratio of fluorescence intensity of NDC80, PRC1, or GFP to that of ACA.

### Protein purification

Bacmid DNA containing GST-NDC80ΔN-FLAG, NUF2ΔN-HA, or GST-PRC1-His was transfected into Sf9 insect cells to produce baculovirus. For protein expression, baculovirus-infected cells were grown at 28°C for 48 hours. After washing with PBS, the cells were stored at −80°C. For protein purification, cells were suspended in 50 mM HEPES (pH 7.4), 300 mM NaCl, 1 mM EDTA, 10% glycerol, 1% Triton X-100, 1 mM DTT and protease inhibitor cocktail (cOmplete EDTA-free, Roche) and lysed by sonication on ice. After centrifugation at 14,000 rpm for 60 minutes, the soluble fraction of the cell lysate was bound to glutathione-sepharose 4B (GST SpinTrap, Cytiva). The unbound fraction was washed with 50 mM HEPES (pH 7.4), 250 mM NaCl, 1 mM EDTA, 10% glycerol and 1 mM DTT. Proteins were eluted by cleavage with PreScission protease (Cytiva), which cleaves GST in 50 mM HEPES (pH 7.4), 150 mM NaCl, 1 mM EDTA, 0.05% Triton X-100 and 1 mM DTT.

### *In vitro* kinase reaction

Purified proteins were incubated with 200 ng human MPS1 (TTK, 05-169, Carna Biosciences) in 16 mM KCl, 32 mM Na-β-glycerophosphate, 8 mM EGTA, 6 mM MgCl_2_, 0.4 mM DTT and 2.5 mM ATP at 37°C for 1 hour. To detect phosphorylation, 1 mM adenosine 5’-O-(3-thiotriphosphate) (ATPγS, ab138911, abcam) was used instead of ATP for the kinase reaction. After kinase reaction, p-nitrobenzyl mesylate (PNBM, ab138910, abcam) was added (final 2.5 mM) and incubated for 1 hour at room temperature.

### Western blotting

Proteins were prepared in an SDS-PAGE sample buffer. After heating at 95°C for 5 minutes, they were detected by Western blotting. The primary antibodies were a mouse anti-FLAG antibody (F1804, Sigma-Aldrich), a rat anti-HA antibody (11867423001, Roche), and a rabbit anti-thiophosphate ester antibody (ab92570, abcam) (1:2000).

Horseradish peroxidase-conjugated anti-mouse, anti-rat, and anti-rabbit antibodies (1:2000) were used as secondary antibodies.

### Mass spectrometry (MS)

Proteins were subjected to SDS-PAGE electrophoresis. Each gel slice underwent in-gel digestion for protein extraction. The slices were diced into 1 mm pieces and then treated with 10 mM tris(2-carboxyethyl)phosphine hydrochloride (SIGMA) at 56°C for 30 minutes for reduction, followed by alkylation with 55 mM iodoacetamide at room temperature for 45 minutes in the dark. Subsequently, digestion was carried out using Trypsin (MS Grade, Thermo Scientific) at 37°C for 16 hours. The resulting peptides were extracted using 1% trifluoroacetic acid and 50% acetonitrile. Phosphorylated peptides were enriched either using the High-Select Fe-NTA Phosphopeptide Enrichment Kit (Thermo Scientific) or the Titansphere Phos-TiO Kit (GL Science) following the manufacturer’s instructions. Both the trypsin digest before enrichment and the enriched fractions were desalted using in-house C18 stage-tip and then used for LC-MS/MS analysis.

Mass spectra were acquired using a Thermo Scientific LTQ-Orbitrap Velos Pro connected to a nano-flow UHPLC system (ADVANCE UHPLC; AMR Inc.) with an Advanced Captive Spray SOURCE (AMR Inc.). Peptide mixtures were injected onto a C18 trap column (PepMap Neo Trap Cartridge, ID 0.3 mm x 5 mm, particle size 5 μm, Thermo Fisher Scientific) and subsequently fractionated by C18 reverse-phase chromatography (3 μm, ID 0.075 mm × 150 mm, CERI). Peptides were eluted with a linear gradient of solvent B (5– 35% acetonitrile, 0.1% formic acid) at a flow rate of 300 nL/min over 60 minutes. The mass spectrometer performed seven successive scans, starting with a full MS scan from 350 to 1600 m/z using Orbitrap (resolution = 60,000), followed by data-dependent scans of the top three most abundant ions using CID in the ion trap for the second to fourth scans, and using HCD in the Orbitrap (resolution = 7,500) for the fifth to seventh scans. Automatic MS/MS spectra were acquired from the highest peak in each scan, with a relative collision energy set to 35% CID or HCD and an exclusion time of 90 seconds for ions within the same m/z range.

The raw files were searched against the Uniprot *Mus musculus* proteome database (downloaded January 2022) and cRAP contaminant proteins dataset using the MASCOT program (version 2.6; Matrix Science) via Proteome Discoverer 2.5 (Thermo Fisher Scientific). The search was conducted with carbamidomethylation of cysteine as a fixed modification, and oxidation of methionine, acetylation of protein N-termini, and phosphorylation of serine, threonine, and tyrosine as variable modifications. The number of missed cleavage sites was set to 2.

### Statistical analysis

Graphing and statistical analysis were performed using Excel, R, and GraghPad Prism. Statistical tests used were described in figure legends.

## Acknowledgements

We thank R. Benezra for providing the NDC80 antibody; J. Ellenberg for providing macros for automated microscopy; and the imaging, genome analysis, and animal facilities of RIKEN Kobe for technical support. We also thank our laboratory members for discussion. This work was supported by RIKEN intramural grants, JSPS KAKENHI JP18H05549/JP21H02407/23H04948 to T.S.K., and JSPS KAKENHI JP17K15069/JP19K06682/JP22K06257 to S.Y.

## Author contributions

S.Y., and T.S.K. conceived the project. S.Y. performed almost all the experiments. R.N. performed mass spectrometry analysis. T.S.K. supervised the project. S.Y., R.N, and T.S.K. wrote the manuscript.

---

**Figure S1.**
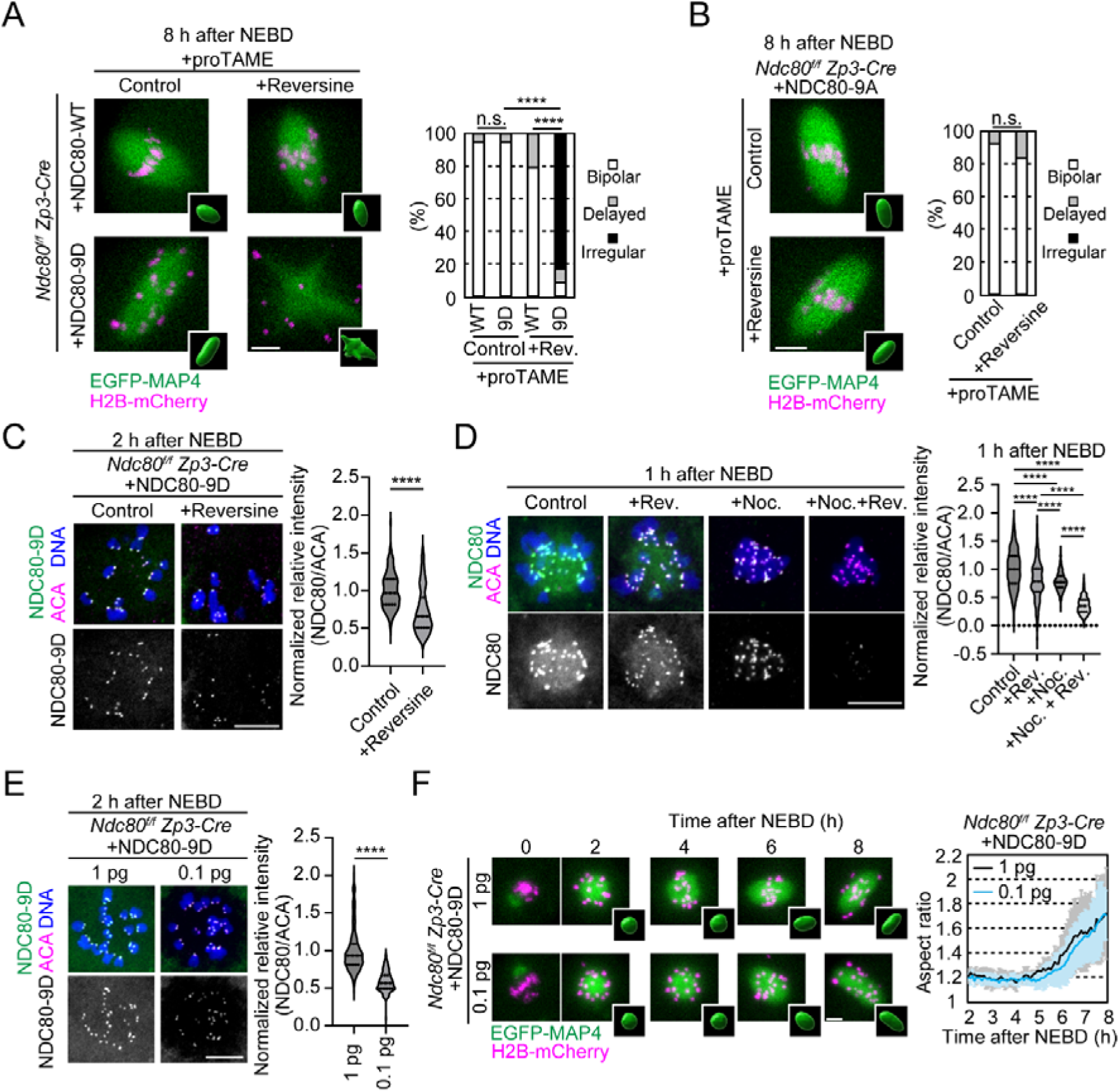
MPS1 activity is required for acentrosomal spindle bipolarization in the absence of stable kinetochore-microtubule attachment. **A.** MPS1 inhibition impairs spindle bipolarization in NDC80-9D oocytes even after metaphase arrest. Live imaging of *Ndc80^f/f^ Zp3-Cre* oocytes expressing EGFP-MAP4 (spindle, green), H2B-mCherry (chromosome, magenta) and NDC80-WT/-9D in the presence of proTAME. Z-projection and 3D-reconstruction images at 8 hours after NEBD are shown. Spindle morphology classification is shown (n=20, 20, 24, 24 oocytes from 3 independent experiments). n.s., not significant, ****p<0.0001 by Fisher’s exact test for “bipolar” groups with Holm correction. **B.** MPS1 inhibition does not impair spindle bipolarization in NDC80-9A oocytes. *Ndc80^f/f^ Zp3-Cre* oocytes expressing NDC80-9A were analyzed as in A (n=26, 24 oocytes from 3 independent experiments). **C.** MPS1 inhibition decreases NDC80-9D localization at kinetochores. Immunostaining of NDC80-9D-GFP, ACA (kinetochores), and Hoechst33342 (DNA) in oocytes treated with reversine. Normalized relative intensities of NDC80-9D-GFP are shown (median and quartiles, n=280, 174 kinetochores from 7, 7 oocytes. Three independent experiments were performed). ****p<0.0001 by two-tailed unpaired Student’s t-test. **D.** MPS1 inhibition decreases NDC80 localization at kinetochores. Immunostaining of anti-NDC80, ACA (kinetochores), and Hoechst33342 (DNA) in oocytes treated with reversine and/or nocodazole at 1 hour after NEBD. Normalized relative intensities of NDC80 are shown (median and quartiles, n=240, 240, 240, 232 kinetochores from 6, 6, 6, 6 oocytes. Three independent experiments were performed.). ****p<0.0001 by Tukey’s multiple comparison test. **E.** Titration of NDC80-9D. Immunostaining of NDC80-9D-GFP, ACA (kinetochores) and Hoechst33342 (DNA) in oocytes expressing 1 pg or 0.1 pg mRNA of NDC80-9D-GFP. Normalized relative intensities of NDC80-9D-GFP are shown (median and quartiles, n=200, 200 kinetochores of 5, 5 oocytes from one experiment). ****p<0.0001 by two-tailed unpaired Student’s t-test. **F.** Reduced NDC80-9D can support spindle bipolarization. Live imaging of *Ndc80^f/f^ Zp3-Cre* oocytes expressing EGFP-MAP4 (spindle, green), H2B-mCherry (chromosome, magenta), and NDC80-9D. Temporal changes in the aspect ratio of the spindle are shown (mean ± SD, n=7, 6 oocytes from one experiment). Scale bars, 10 μm.

**Figure S2.**
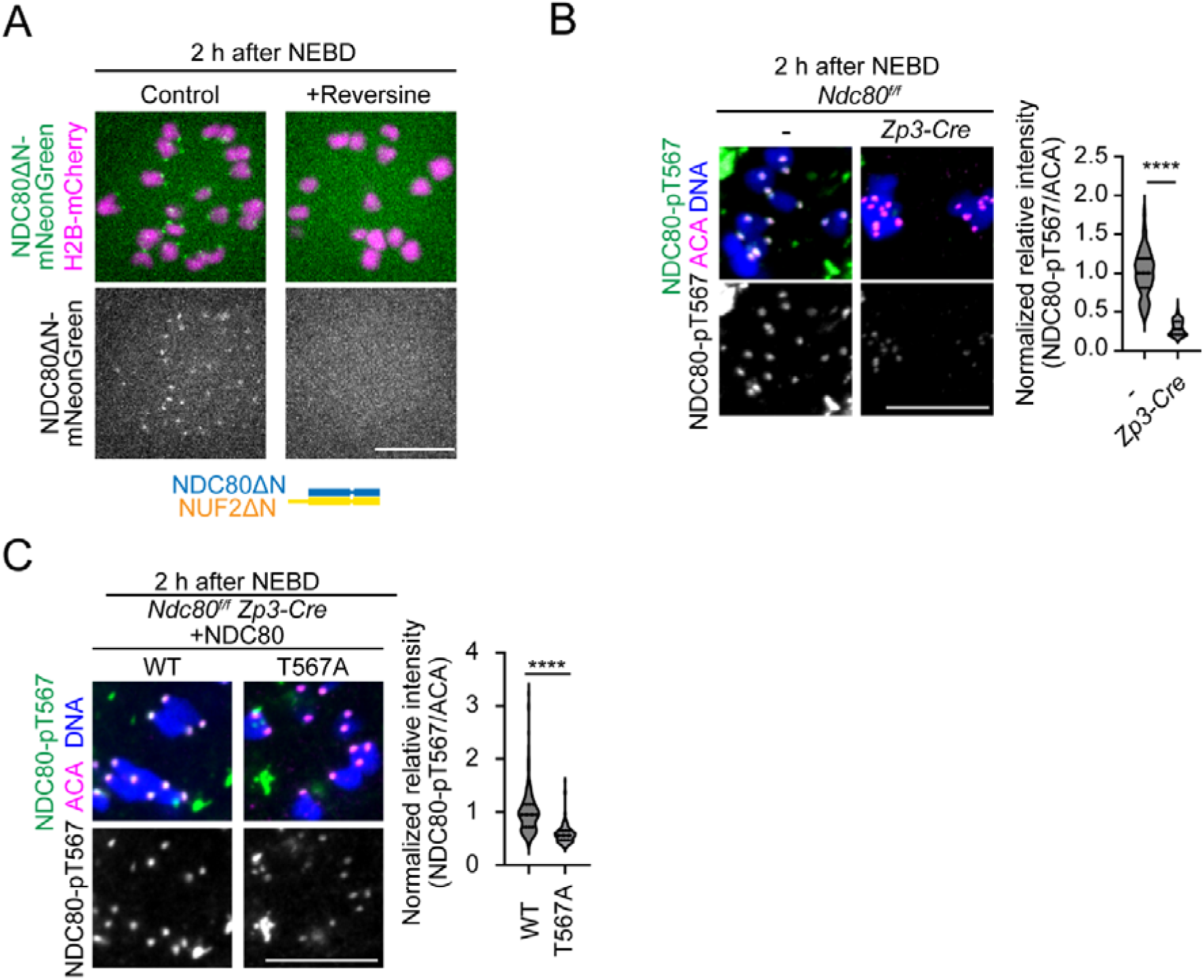
MPS1 promotes spindle bipolarization via the C-terminal domains of NDC80-NUF2 during prometaphase. **A.** MPS1 inhibition delocalizes NDC80ΔN from kinetochores. *Ndc80^f/f^ Zp3-Cre* oocytes expressing NDC80ΔN-mNeonGreen, NUF2ΔN, and H2B-mCherry treated with reversine were imaged. **B.** Phospho-antibody specificity. *Ndc80^f/f^ Zp3-Cre* oocytes were immunostained with anti-phospho-NDC80-T567 antibody, ACA (kinetochores), and Hoechst33342 (DNA). Control oocytes were without *Zp3-Cre.* Normalized relative intensities of phospho-NDC80-T567 are shown (median and quartiles, n=199, 168 kinetochores from 5, 5 oocytes. Three independent experiments were performed). ****p<0.0001 by two-tailed unpaired Student’s t-test. **C.** Phospho-antibody specificity. *Ndc80^f/f^ Zp3-Cre* oocytes expressing NDC80-WT/-T567A were immunostained with anti-phospho-NDC80-T567, ACA (kinetochores), and Hoechst33342 (DNA). Normalized relative intensities of NDC80-pT567 are shown (median and quartiles, n=240, 240 kinetochores from 6, 6 oocytes. Three independent experiments were performed). ****p<0.0001 by two-tailed unpaired Student’s t-test. Scale bars, 10 μm.

**Figure S3.**
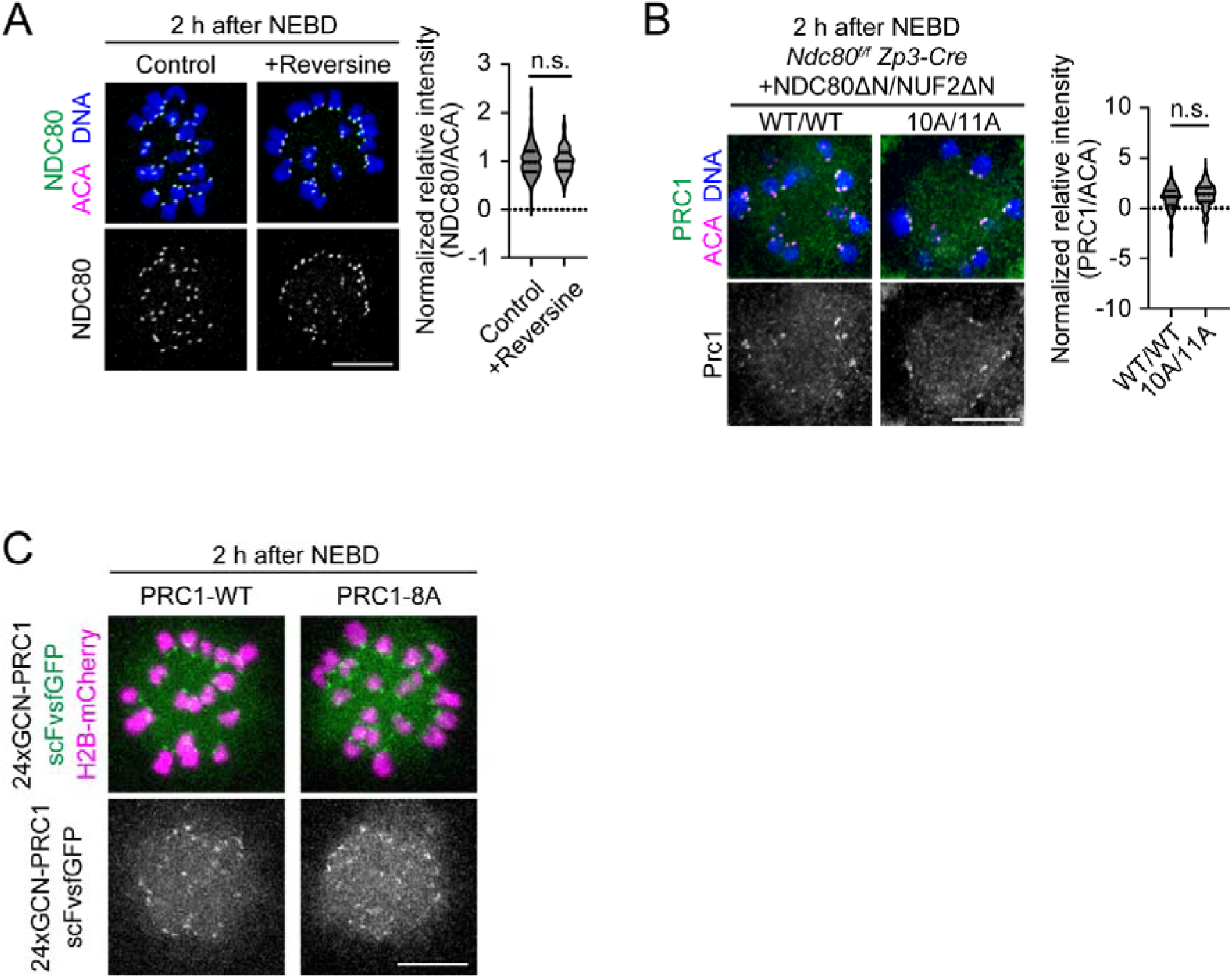
MPS1 activity promotes kinetochore localization and spindle bipolarization activity of PRC1. **A.** NDC80 is not significantly decreased in MPS1-inhibited oocytes at 2 hours after NEBD. Immunostaining of reversine-treated oocytes with anti-NDC80, ACA (kinetochores), and Hoechst33342 (DNA). Normalized relative intensities of NDC80 are shown (median and quartiles, n=200, 200 kinetochores from 5, 5 oocytes. Three independent experiments were performed.). n.s., not significant by two-tailed unpaired Student’s t-test. **B.** PRC1 can be recruited to kinetochores by phospho-mutant NDC80ΔN and NUF2ΔN. Immunostaining of *Ndc80^f/f^ Zp3-Cre* oocytes expressing NDC80ΔN-WT/-10A and NUF2ΔN-WT/-11A with anti-PRC1, ACA (kinetochores), and Hoechst33342 (DNA). Normalized relative intensities of PRC1 are shown (median and quartiles, n=200, 200 kinetochores from 5, 5 oocytes. Three independent experiments were performed.). n.s., by two-tailed unpaired Student’s t-test. **C.** Phospho-mutant PRC1 can localize to kinetochores. Oocytes expressing 24xGCN-PRC1-WT/-8A, scFvsfGFP, and H2B-mCherry were imaged. Scale bars, 10 μm.

